# Long Reads Capture Simultaneous Enhancer-Promoter Methylation Status for Cell-type Deconvolution

**DOI:** 10.1101/2021.01.28.428654

**Authors:** Sapir Margalit, Yotam Abramson, Hila Sharim, Zohar Manber, Surajit Bhattacharya, Yi-Wen Chen, Eric Vilain, Hayk Barseghyan, Ran Elkon, Roded Sharan, Yuval Ebenstein

## Abstract

**Motivation:** While promoter methylation is associated with reinforcing fundamental tissue identities, the methylation status of distant enhancers was shown by genome-wide association studies to be a powerful determinant of cell-state and cancer. With recent availability of long-reads that report on the methylation status of enhancer-promoter pairs on the same molecule, we hypothesized that probing these pairs on the single-molecule level may serve the basis for detection of rare cancerous transformations in a given cell population. We explore various analysis approaches for deconvolving cell-type mixtures based on their genome-wide enhancer-promoter methylation profiles.

**Results:** To evaluate our hypothesis we examine long-read optical methylome data for the GM12787 cell line and myoblast cell lines from two donors. We identified over 100,000 enhancer-promoter pairs that co-exist on at least 30 individual DNA molecules per pair. We developed a detailed methodology for mixture deconvolution and applied it to estimate the proportional cell compositions in synthetic mixtures based on analyzing their enhancer-promoter pairwise methylation. We found our methodology to lead to very accurate estimates, outperforming our promoter-based deconvolutions. Moreover, we show that it can be generalized from deconvolving different cell types to subtle scenarios where one wishes to deconvolve different cell populations of the same cell-type.

**Availability:** The code used in this work to analyze single-molecule Bionano Genomics optical maps is available via the GitHub repository https://github.com/ebensteinLab/Single_molecule_methylation_in_EP.

**Contact:** (Y.E), (R.S)

## 1 Introduction

The accumulation of high-throughput genome-wide methylation data has enabled the analysis of human methylomes across distinct populations and medical cohorts via epigenome-wide association studies (EWAS) (Gorenjak *et al*., 2020; Küpers *et al*., 2019; Chu *et al*.). Such analyses have shown that predisposition to common human disease is frequently associated with specific methylation signatures in distal control regions also known as gene enhancers (Li *et al*., 2013). While the contribution of DNA methylation in gene promoters to variation in intertumor gene expression was found to be low, enhancer methylation provides a much higher level of contribution to tumor heterogeneity and may further illuminate the mechanism of cancer predisposition (Li *et al*., 2013). Furthermore, changes in DNA methylation patterns have been shown to correlate with early carcinogenesis, even prior to tumor formation, as well as with metastasis and response to therapy (Kurkjian *et al*., 2008; Hentze *et al*., 2019; Vrba and Futscher, 2020). Aran *et. al*. have shown that enhancer methylation is drastically altered in cancers and is closely related to altered expression profiles of cancer genes (Aran and Hellman, 2013a, 2013b; Aran *et al*., 2013). Hansen *et.al*. have shown that regions which are differentially methylated between cancer and normal tissue are more prone to variability in methylation levels, suggesting that stochastic epigenetic variation is a fundamental characteristic of the cancer phenotype (Hansen *et al*., 2011). Nevertheless, available analyses only assess enhancer-promoter methylation on the population level, averaging out any differences between individual cells in the studied sample. We hypothesize that variability in methylation on the population level may be attributed to variability in the mixing ratios of cancer and benign phenotypes of the same cell type. In such cases the detailed single-cell enhancer-promoter methylation profile may provide valuable information for studying the evolution of early carcinogenesis and tumor heterogeneity. Long reads present a unique opportunity to study the co-existence of methylation in a promoter and its enhancers along the same DNA molecule, in effect providing single-cell information for the studied locus. When examining many such molecules, the methylation pattern distribution of an enhancer-promoter pair may be directly recorded and used to enumerate cell subsets in a mixture, similar to what may be achieved with gene expression RNA mixtures (Newman *et al*., 2015; Zaitsev *et al*., 2019). Methylation profiles have already been utilized to infer cell mixture distribution with good accuracy (Houseman *et al*., 2012). It remains to be tested whether pairwise analysis of enhancer-promoter pairs provides information on subtle transformations in cells with identical genetic backgrounds such as in early cancer. In order to establish the analytical framework for such data we analyzed whole genome Bionano Genomics optical methylation maps (Sharim *et al*., 2019). We identified ~4 million long molecule reads encompassing enhancer-promoter pairs up to 200kb apart at 30X-300X coverage and explored several analytical approaches to harness these data for cell mixture deconvolution.

## 2 Methods

### 2.1 Data collection

Single-molecule methylation maps of three replicates of the B-lymphocyte cell line GM12878 were adapted from (Sharim *et al*., 2019), and similar maps of immortalized myoblasts of two human subjects (Coriell) were obtained by the same methods. Shortly, high molecular weight DNA was extracted to ensure long single molecules. DNA was then fluorescently barcoded at specific sequence motifs for alignment to an in-silico reference. Unmethylated cytosines in the recognition sequence TCGA were fluorescently labeled to perform reduced representation optical methylation mapping (ROM) on the Bionano Genomics Saphyr instrument. The genomic location of the labeled unmodified cytosines on individual DNA molecules was inferred from direct alignments to the hg38 reference (Bionano Access and Solve). Label coordinates were extended to 1 kb to account for the optical measurement resolution, limiting localization accuracy to ~1000 bp (Wang *et al*., 2012).

### 2.2 Distal genomic elements links and coordinates

Predicted enhancers were mapped to genes by the JEME engine (Cao *et al*., 2017). Genomic coordinates of enhancers were converted from the human genome build hg19 to hg38 using UCSC liftOver (Haeussler *et al*., 2019)). Ambiguous genomic regions (Amemiya *et al*., 2019) were subtracted from enhancer locations, and enhancers smaller than 200 bp were extended to 200 bp around their midpoint. Promoters were defined according to the transcription start sites (TSS) of the protein-coding genes (Gencode V.34 annotations, (Frankish *et al*., 2019)), and taken as 2000 bp upstream and 500 bp downstream of the TSS. Enhancers and promoters not containing at least one potential site for methylation labeling were discarded. E-P pairs in close proximity, less than 5,000 bp, were also filtered out. All predicted enhancers under these conditions were used for our analysis, regardless of cell type and biological contexts used for prediction. For comparison against promoter-only analysis, we also created a dataset with a single enhancer assigned to each promoter. We focused on enhancers that display the highest number of potential detectable methylation sites. In case of tie, enhancer size and proximity to the corresponding promoter were also considered. These criteria are unbiased toward specific cell types and select enhancers with the highest potential for reliable methylation calling by ROM.

### 2.3 Coexistence of methylation signals at the distal elements

We focused on DNA molecules that span entire enhancer-promoter pairs (E-P) and recorded the corresponding methylation states. The enhancers and promoters’ states were reduced to binary methylation states: if the element showed any degree of fluorescence, it was identified as ‘unmethylated’. Therefore, every enhancer and promoter pair coexisting on a DNA molecule displays one of four possible methylation combinations with the promoter and enhancer being methylated or unmethylated. For every pair, the number of molecules belonging to each class is counted in order to record the exact pairwise methylation distribution. Enhancer-promoter pairs that were covered by less than 30 molecules in an experiment were filtered out.

### 2.4 Matched promoter methylation data

To benchmark our pairwise analysis we extracted a dataset of promoter only methylation from the same molecules that span the E-P pairs. These were used to assess if the coexistence of enhancer-promoter pairs holds similar or additional information beyond traditional promoter methylation analysis.

### 2.5 Assembling mixtures and training/test division of molecules

To test our deconvolution pipeline, two different mixtures were simulated from three replicates of the B-lymphocyte cell line GM12878 and two lines of immortalized myoblast cells from different donors:

1. Mixture of different cell types – B-lymphocytes and myoblasts: Molecules of the three GM12878 replicates were merged to a single experiment, and molecules of the two myoblasts were merged as well.
2. Mixture of the same cell type from different individuals: myoblast cells from two different human subjects.

In each set, molecules of the two different experiments (referred to as sample “A” and sample “B”) were randomly divided between a training set, that was kept pure, to be used in the supervised deconvolution of mixtures, and a test set, that was mixed in known ratios with the test set molecules of the other experiment (0-100% in respect to one of the samples, in 10% increments). Test sets of both samples contained an equal number of molecules, accounting for 33% of molecules in the smaller experiment.

### 2.6 Deconvolution of mixtures

We explored several methods for deconvolving the mixtures to infer the mixing ratio: (i) local projection of vectors, (ii) global minimization of sum of squared errors (SSE) and Kullback-Leibler divergence (KLD) measures, and (iii) global maximum likelihood estimation (MLE). We describe these methods in detail below.

#### 2.6.1 Vector projection

In this local method, the mixing ratio is calculated separately for each enhancer-promoter pair. Specifically, the normalized proportions of each of the four possible methylation combinations define a 4-dimensional vector representing the pair in a given sample. Each E-P pair is characterized by three such vectors belonging to the test set 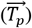 and the two pure training sets 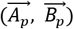. In order to assess the proportion of molecules in the test set originating from each pure sample, the difference between the test set vector and the vector 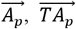, was projected on the difference between vectors 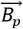 and 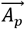 of the training sets, 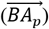. The mixing ratio w.r.t. sample B is given by the ratio between the size of the difference between the projection vector and vector 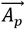 (calculated as the dot product of 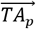 and 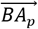 divided by the size of 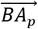), and the size of vector 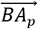. The mixing ratio with respect to sample A completes this value to 1. The final selected mixing ratio is the average of mixing values calculated for all E-P pairs (Eq. 1).

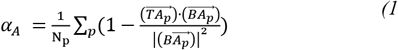

*α_A_*, is the mixing ratio relative to sample A. *p* represents the E-P pair. N_p_ is the total number of E-P pairs. 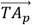 represents the difference between the test set vector and the vector of the training set of sample A; 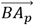 represent the difference between the vectors of training sets B and A.

#### 2.6.2 Minimizing the difference between the test set and linear combinations of the training set via SSE and KLD computations

The counts of the different methylation combinations in each enhancer-promoter pair in the mixed test set and the pure training sets were normalized to obtain ratios, making sure all ratios are non-zeros by adding 0.01 to each ratio and renormalizing to 1. The ratios of the pure training sets were treated as the probability of a molecule spanning a certain E-P pair to have a specific methylation combination given the sample it came from, either “A” or “B”. Then, linear combinations of the two pure training sets were assembled per pair in all possible mixing ratios w.r.t. sample A (0-100% in 1% increments).

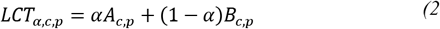

*LCT_α,c,p_* is the linear combination of the pure training sets per EP pair (*p*), methylation combination (*c*), and current parameter of the training sets linear combination (*α*) *A_c,p_* and *B_c,p_* are the probabilities of molecules from training sets A and B respectively to have methylation combination *c* given the sample and the E-P pair *p*.

The test set is compared against all the 101 linear combinations of the training sets. The ratio *α* that minimizes these expressions is reported. We use two minimization criteria:

##### 2.6.2.1 Sum of squared errors (SSE)

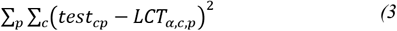

*p* represents the E-P pairs, *c* represents methylation combinations, *α* represents the current parameter of the training sets linear combination.

##### 2.6.2.2 Kullback-Leibler divergence (KLD)

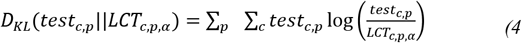

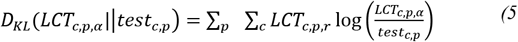

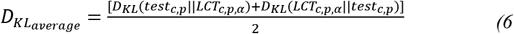

*p* represents the E-P pairs, *c* represents methylation combinations, *α* represents the current parameter of the training sets linear combination.

#### 2.6.3 Maximum likelihood estimation (MLE)

Last, we considered a method that is based on a probabilistic model of the data which asserts that each molecule is chosen from one of the samples based on the mixing ratio and then its methylation status is chosen based on the corresponding normalized counts vector. In detail, the probability of a molecule in the test set to originate from sample A, is the mixing ratio, α, whereas its probability to originate from sample B is 1-α (Eq. 7).

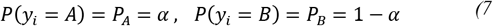

*i* stands for a molecule in the test set, y is the hidden information about the set it came from.

The probability of a molecule from the test set to exhibit a methylation combination *c*, given the sample it came from (A or B) and the E-P pair it spans can be evaluated as the proportion of combination *c* in that E-P pair in the training set of the corresponding sample (Eq. 8).

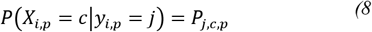

X_i,p_ is the observed methylation combination of molecule *i* of pair *p*; *j* is the sample of origin, either A or B.

We estimate the mixing ratio for which the test data observations are most probable by maximizing the log-likelihood function (Eq. 9):

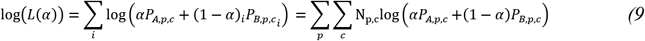

Where *i* a test set molecule, *p* is the pair it belongs to, *c* is its methylation combination, A and B are the two samples, *N_p,c_* is the number of molecules in the test set that come from pair *p* and display methylation combination *c*.

As the log likelihood is concave, we can find its maximum using gradient ascent. In our implementation of the algorithm (Eq. 10), the difference in log-likelihood from previous iteration served as a stopping criterion (0.001), while confining the mixing ratio to the relevant range (0-1), as well as limiting the number of iterations (no more than 20,000). The mixing ratio that yielded the highest log-likelihood is reported.

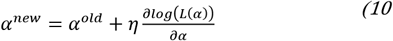

*α^new^* is the calculated mixing value in the current iteration. *α^old^* is the mixing ratio obtained in the previous iteration (initialized to 0.5). *η* is the step size of the gradient ascent algorithm (fixed to 0.005).

### 2.7 Supervised selection of enhancer-promoter pairs

Several methods are proposed here to rank the different E-P pairs by their ability to discriminate between the training set samples. This ranking can serve to select a smaller subset of pairs, to ensure more accurate results and noise filtration. We explored three such methods, as detailed below.

#### 2.7.1 Euclidean distances

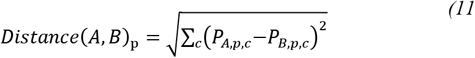

*p* is the current E-P pair, *c* represents the methylation combinations, *P_A,p,c_*, *P_B,p,c_* are the proportions of methylation combination *c* in pair *p* in samples A and B respectively.

#### 2.4.2 KL divergence

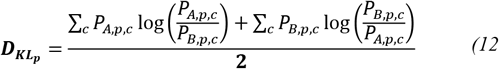

*p* is the current E-P pair, *c* represents the methylation combinations, *P_A,p,c_*, *P_B,p,c_* are the proportions of methylation combination *c* in pair *p* in samples A and B respectively.

#### 2.7.3 Weighted KLD (wKLD)

Molecule coverage of the different enhancer-promoter pairs in each training sample can vary Multiplying the KLD score of each combination of every pair by the number of molecules displaying this combination in the corresponding sample and pair, puts more weight on the highly-covered pairs, which presumably provide more reliable information.

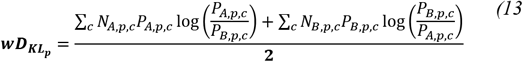

*p* is the *p* is the current E-P pair, *c* represents the methylation combinations, *P_A,p,c_*, *P_B,p,c_* are the proportions of methylation combination *c* in pair *p* in samples A and B respectively. *N_A,p,c_*, *N_B,p,c_* are the number of molecules in pair *p* that display methylation combination *c* in the training sets of samples A and B respectively.

## 3 Results

Our analytical exploration builds on the fact that new types of data that profile genomic methylation *via* long single molecule reads have recently become available. Of special interest are datasets obtained by Bionano Genomics optical genome mapping. These data contain the largest fraction of molecules longer than 100 kbp in comparison to other long read approaches, allowing us to explore distant enhancers in the context of their molecular promoter. The method is inherently poor in resolution but may effectively report on methylation status at the level of genomic elements such as gene bodies, promoters and enhancers. Fig. 1 shows a stack of digitized DNA molecules mapped to a region in chromosome 17 containing the TP53 gene locus and three of its predicted enhancers, located ~50-100 kb away from the promoter. All individual molecules selected from these data span the gene promoter and at least one distant enhancer. Methylation labels shown in dark blue along the grey molecules contour denote unmethylated CpGs (non-methylation labels have been artificially enhanced inside the boxed promoter and enhancer regions). It can be clearly seen that the promoter and the two closest enhancers are highly labelled and thus unmethylated. The leftmost enhancer is almost void of methylation labels, indicating that it is highly methylated. The methylation status is well reflected in the average methylation profile shown on the bottom of the figure and reflecting the accumulation of methylation labels from all molecules along the region. We note that for many gene promoters and enhancers the methylation status is variably distributed along individual molecules, with all four combinations of E-P methylation represented for some genes.

**Fig. 1.**
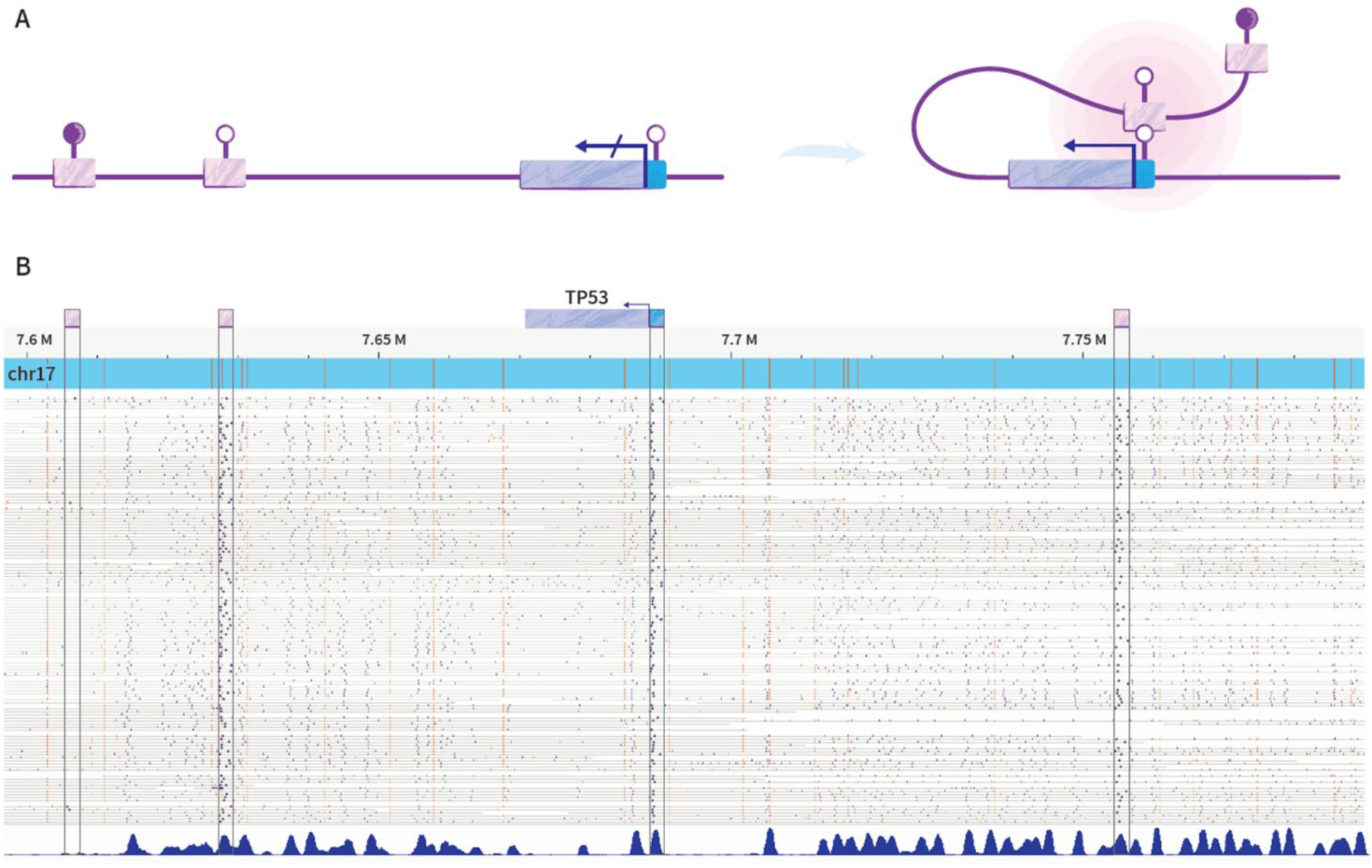
Methylation states in predicted enhancer-promoter pairs. A. schematic illustration of possible methylation states for a promoter and enhancers, and potential interaction between them. B. Bionano genomics optical methylation map of a region in chromosome 17 in GM12878 DNA. The region contains the gene TP53, its promoter (small blue box), and several predicted enhancers (pink boxes). Blue dots denote unmethylated sites and orange dots denote genetic tags used for alignment to the hg38 reference.

### 3.1 Comparison between deconvolution methods for promoters and E-P pairs

We first set out to compare the deconvolution efficacy of E-P pairwise methylation in comparison with promoter methylation. Given that each promoter has multiple predicted enhancers which results in a much larger E-P dataset, we restricted this analysis to a single enhancer assigned to each promoter as described in section 2.2. Deconvolution of simulated mixtures was performed by several methods: local projection of vectors, global minimization of the sum of squared errors (SSE), Kullback-Leibler divergence (KLD), and maximum likelihood estimation (MLE) (see Methods). Mixtures containing two different cell types (B-lymphocytes and myoblasts) at 10% increments were subject to deconvolution by each of the methods for both promoter-only methylation as well as in the context of E-P pairwise methylation. The least accurate deconvolution was achieved by vector projections, yielding over 7% average error for the promoter-based analysis and over 4% error for the E-P analysis. The best deconvolution was achieved by MLE with 0.86% for promoter methylation and 0.69% for E-P methylation (Fig. 2.).

**Fig. 2.**
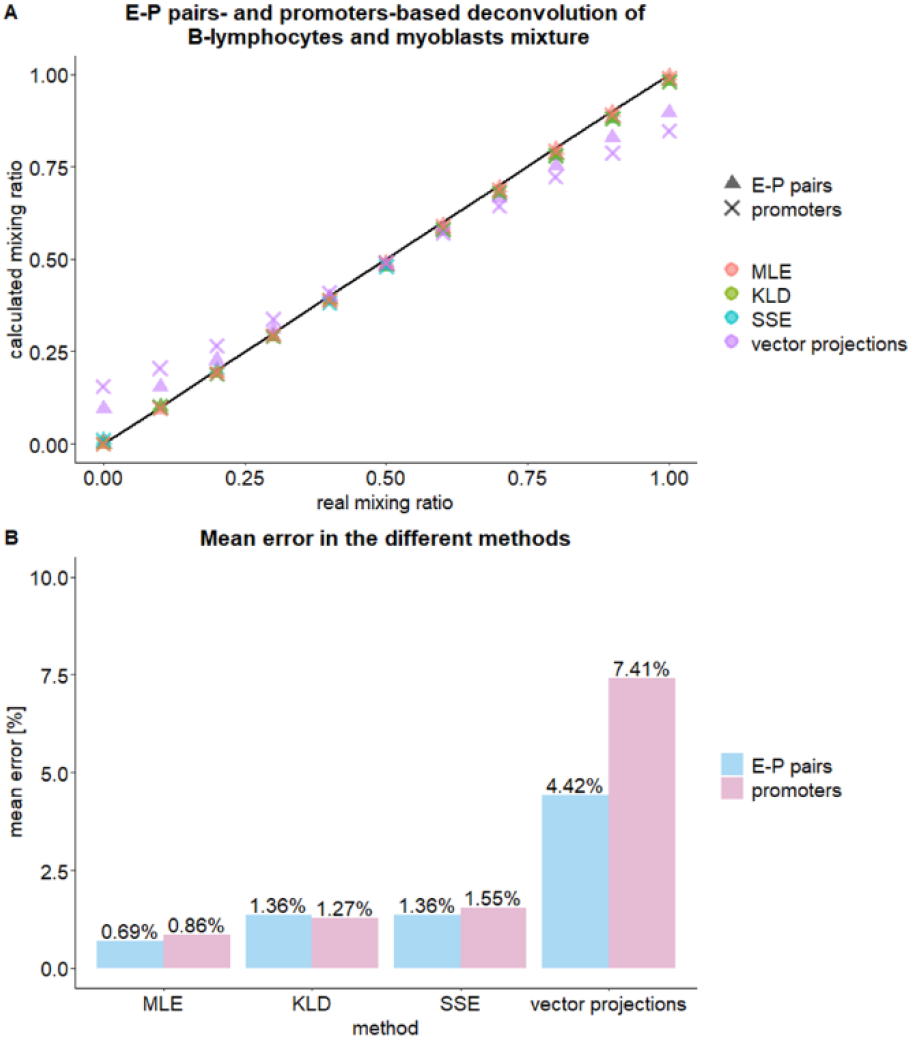
Deconvolution of B-lymphocytes and myoblast cells mixtures by different methods using methylation states in promoters alone and enhancer-promoter pairs, accounting for one enhancer per promoter. A. calculated mixing ratio according to the different methods vs. the known mixing ratio. B. the mean error in calculated mixing ratio, calculated as the absolute distance from the known ratio, in the different methods.

### 3.2 Deconvolution of myoblasts derived from two individuals using full E-P methylation

The abundance of enhancers and the interplay between their interaction with their gene promoters may hold important information on the precise state of a cell. While for comparison with promoter-based analysis we limited the number of enhancers to one per promoter, our E-P dataset is composed of over 100,000 different pairs with detailed pairwise distribution for each pair. B-lymphocytes and myoblasts, two distinct cell types, were successfully resolved with 1.01-1.36% accuracy by three of the four methods tested (Fig. 3a.). Additionally, the incorporation of multiple enhancers per promoter does not show any significant difference in deconvolution performance relative to the more limited set used for comparison with promoters (Fig. 3b.). Nevertheless, analyzing the full E-P methylation dataset is more biologically relevant as it does not make assumptions on the activity of enhancers and inherently contains more information (but also more noise).

**Fig. 3.**
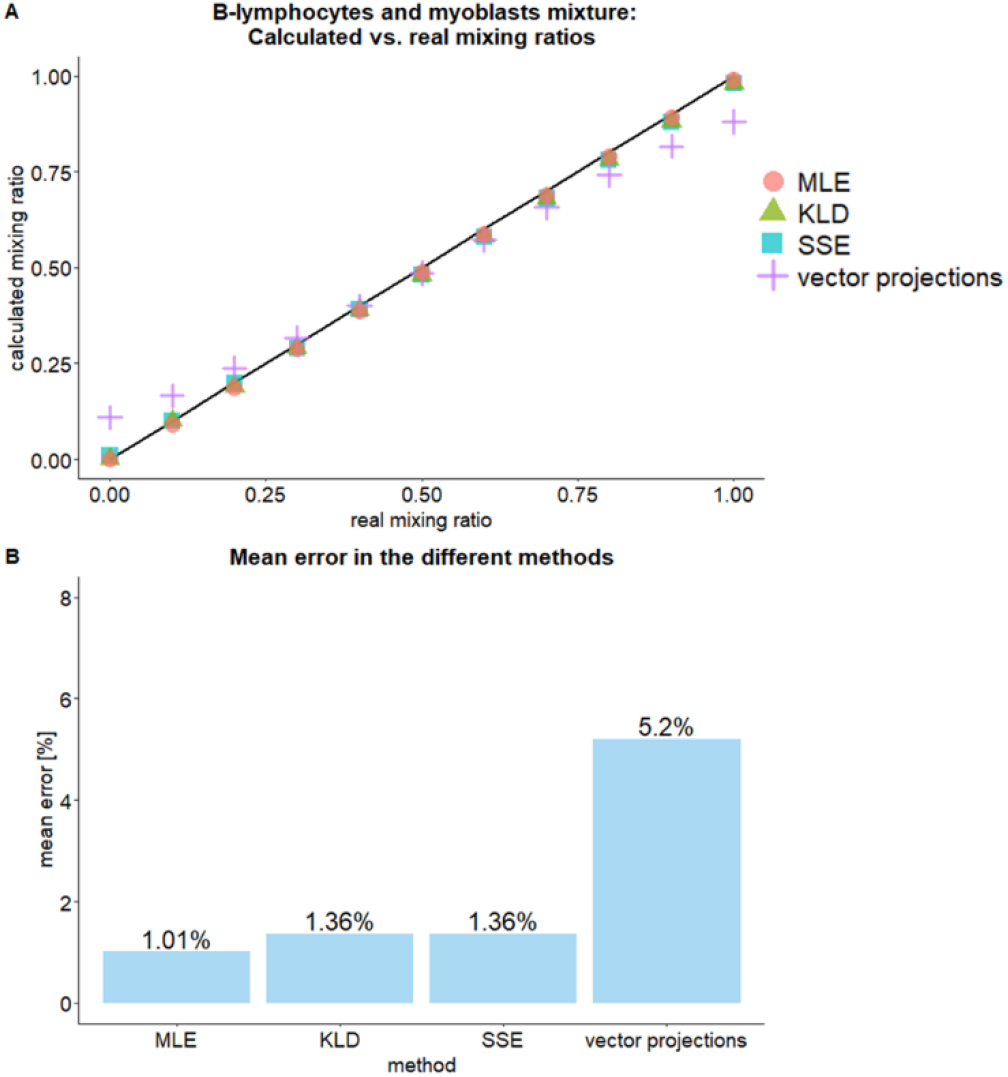
Deconvolution of B-lymphocytes and myoblast cells mixtures by different methods using methylation states in all predicted enhancer-promoter pairs. A. calculated mixing ratio according to the different methods vs. the known mixing ratio. B. the mean error in calculated mixing ratio, calculated as the absolute distance from the known ratio, in the different methods.

We next tested our deconvolution methods on mixtures of two myoblast cell lines from different donors. The mixtures were resolved with 5.81-6.82% accuracy by the same three methods (fig. 4). The two myoblast samples are more similar to each other in genome-wide methylation profiles than they are to the unrelated B-lymphocytes in the previous mixtures. Hence, random alterations between the samples may occupy more weight, and can explain the decline in accuracy. We hypothesized that a smaller, educated subset of E-P pairs used for deconvolution could improve the analysis.

**Fig. 4.**
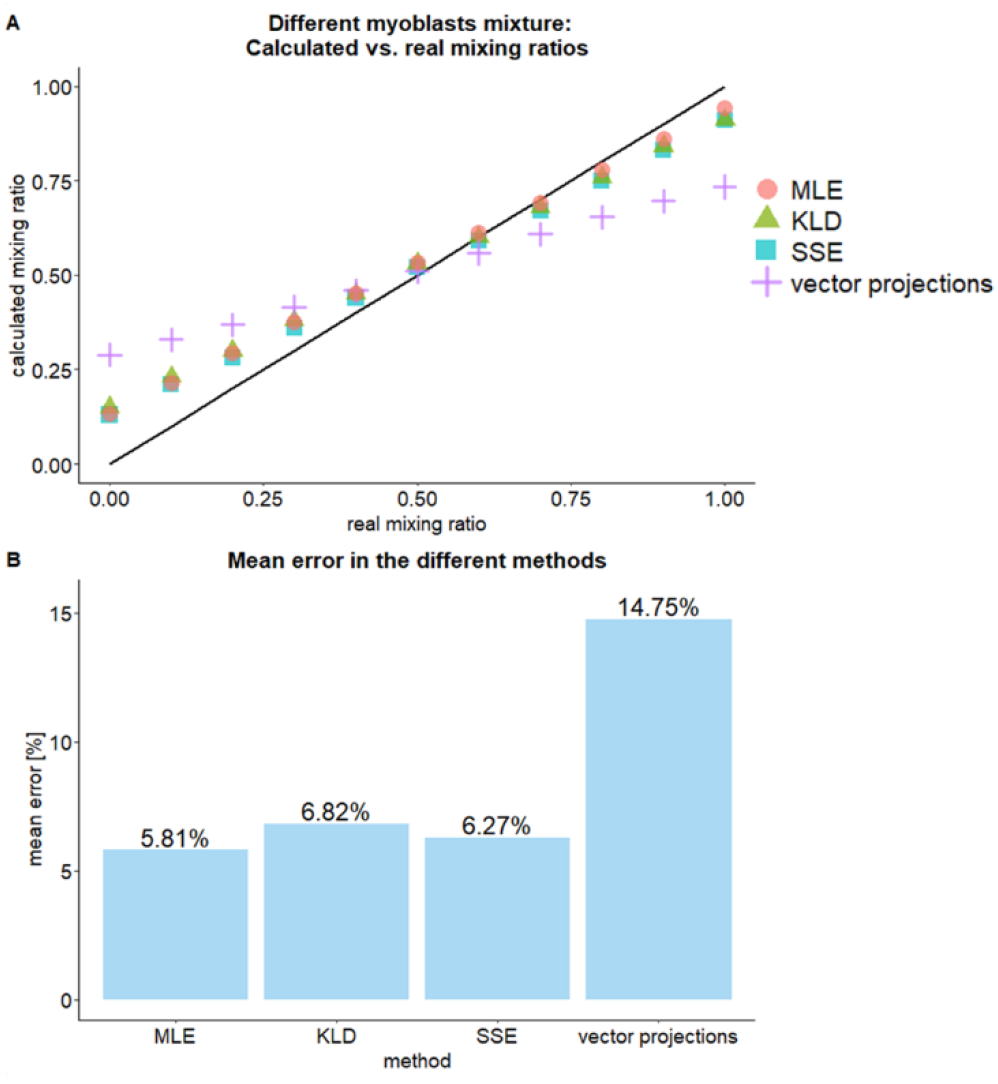
Deconvolution of myoblast cells from two different donors by different methods. A. calculated mixing ratio according to the different methods vs. the known mixing ratio. B. the mean error in calculated mixing ratio, calculated as the absolute distance from the known ratio, in the different methods.

### 3.3 Supervised selection of enhancer-promoter pairs

With over 100,000 predicted enhancer-promoter links used for deconvolution, it is reasonable to assume that not all pairs contribute equally to the discrimination between sample types. Most probably the identity of the most contributing pairs is specific to the types of samples being resolved. Accordingly, relying only on a subset of most differentiating pairs, while filtering out the rest, has potential to improve deconvolution accuracy by filtering out invaluable and possibly noisy data. Additionally, if a small subset of pairs is sufficient for accurate deconvolution it simplifies and shortens the required analysis. We tested three methods for ranking the pairs: Euclidean distances, KLD, and Weighted KLD (wKLD) (see Methods). The performance of the different deconvolution methods was compared for all ranking methods and with different numbers of highest-ranking pairs selected. The full set constituted 108,048 pairs in the mixture of B-lymphocytes and myoblasts, and 135,793 pairs for the two different myoblasts. The mean deconvolution error for several subsets of highest-ranking pairs are shown in log_10_ scale in Figure 5. The different ranking methods provide similar results and the differences in accuracy are mostly attributed to the deconvolution approach used. The lymphocytes and myoblasts mixtures were resolved with ~1% average deconvolution accuracy by MLE, using 75,000-108,048 pairs, and the mixture of the two myoblasts was resolved with ~1.4-1.8% average accuracy using only 100 pairs chosen by KLD or wKLD with MLE deconvolution.

**Fig. 5.**
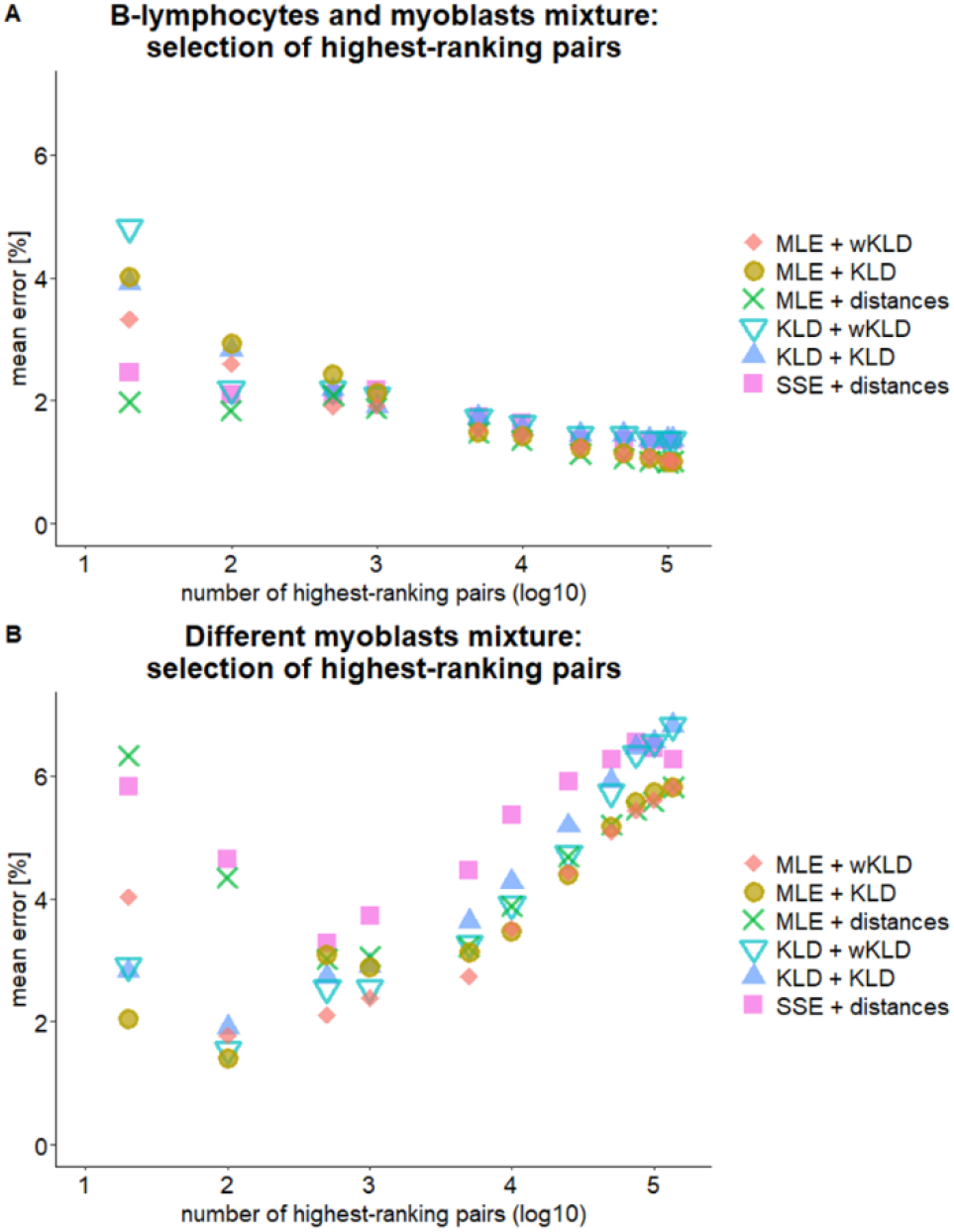
Subsets of E-P pairs, selected by supervised selection methods. The mean error in calculated mixing ratio (distance from theoretical ratio) is displayed against the log10 of the number of best pairs selected in each combination of deconvolution method and ranking method. A) A mixture of two cell types: B-lymphocytes and myoblasts. B) A mixture of two myoblast cells derived from different individuals.

Figure 5 reveals opposite trends in respect to the optimal size of pairs subset used for deconvolution. Whereas the mixture of the different cell types is monotonically resolved more accurately with increasing number of pairs (>75,000 pairs), the mixture of myoblast cells derived from different individuals shows a distinct minimum in the mean error for 100-500 pairs. Since DNA methylation patterns are known to regulate the expression of cell-type specific genes (Dor and Cedar, 2018), a higher variance in methylation signatures is expected between different cell types such as lymphocytes and myoblasts. Different cell-specific methylation patterns imply that more regions along the genome are differential and may contribute to deconvolution, making their differentiation simple and accurate. Deconvolving mixtures of the same cell types such as the mixture of myoblasts from the different individuals is more challenging. We postulate that as may frequently happen in diseased tissue, the observed methylation differences are not related to the cell’s identity, but factors as disease, age, or exposure to environmental stimuli. In such cases methylation variability at cell-type specific loci adds noise to the deconvolution analysis. Sorting the pairs by their information contribution provides a supervised educated approach for assembling the list of pairs that yields the most accurate differentiation.

## 4 Conclusions

This work lays the ground for cell-type deconvolution utilizing a new type of data structure now available *via* long single-molecule methylation maps. This data structure contains chromosome-level methylation profiles of gene bodies, promoters and one or more distant enhancers, all on the same molecule. We test several deconvolution methods and show that for differentiating two cell types, the pairwise analysis yields better deconvolution than promoter-based analysis, reaching an error rate of 0.7%. Since enhancer methylation is known to be a major contributor to methylation variability within a cell-type population such as in cancer, we also analyzed mixtures of two myoblast cell-lines derived from two individuals. The full E-P pair dataset yielded a deconvolution error of ~6% for these highly similar samples. We reasoned that cells with similar methylomes will be differentially methylated only at a subset of loci while variability in common methylation loci will add noise to the deconvolution process. We tested several feature selection algorithms to rank the pairs according to their differentiation capacity. We assessed deconvolution fidelity for various numbers of highest-ranking pairs and found that for the two distinct cell types the deconvolution error monotonically declines with additional pairs. For the two myoblast samples on the other hand, a clear minimum was calculated at ~100 pairs that reduced the error from ~6% to ~1.5%. These results constitute a first step towards harnessing enhancer-promoter linked methylation for deconvolution of cell populations with highly similar cell-type methylomes. Despite focusing on Bionano Genomics reduced-representation optical methylation mapping (ROM), which currently provides the highest coverage of long reads, the principles are valid to other future datasets such as those produced by Oxford Nanopore ultralong-read sequencing protocol. Further exploration of these linkages, including the joint effects of multiple enhancers per promoter may shed light on insightful cellular transformations regulated by long range epigenetic interactions.

## Data availability

The data underlying this article are available on Dropbox via https://www.dropbox.com/sh/kr3ezyljlrjcjxp/AABZVqSsuaHdWZYcJlxKnhWIa?dl=0.

## Acknowledgements

Authors contributions: Y.E and R.S conceived and supervised the project. S.M, Y.A, H.S and Z.M performed data analysis. S.B, Y.W.C, E.V and H.B collected data. R.E advised the project. Y.E, R.S and S.M wrote the manuscript. All authors read and edited the manuscript.

